# Desiccation stress acts as cause as well as cost of dispersal in *Drosophila melanogaster*

**DOI:** 10.1101/2021.03.27.437318

**Authors:** Abhishek Mishra, Sudipta Tung, V.R. Shree Sruti, P.M. Shreenidhi, Sutirth Dey

## Abstract

1. Environmental stress is one of the important causes of biological dispersal. At the same time, the process of dispersal itself can incur and/or increase susceptibility to stress for the dispersing individuals. Therefore, in principle, stress can serve as both a cause and a cost of dispersal.
2. Desiccation stress is an environmentally relevant stress faced by many organisms, known to shape their population dynamics and distribution. However, the potentially contrasting roles of desiccation stress as a cause and a cost of dispersal have not been investigated. Furthermore, while desiccation stress often affects organisms in a sex-biased manner, it is not known whether the desiccation-dispersal relationship varies between males and females.
3. We studied the role of desiccation stress as a cause and cost of dispersal in a series of experiments using *D. melanogaster* adults in two-patch dispersal setups. We were interested in knowing whether (a) dispersers are the individuals that are more susceptible to desiccation stress, (b) dispersers pay a cost in terms of reduced resistance to desiccation stress, (c) dispersal evolution alters the desiccation cost of dispersal, and (d) females pay a reproductive cost of dispersal. For this, we modulated the degree of desiccation stress faced by the flies as well as the provision of rest following a dispersal event.
4. Our data showed that desiccation stress served as a significant cause of dispersal in both sexes. Further investigation revealed an increase in both male and female dispersal propensity with increasing desiccation duration. Next, we found a male-biased cost of dispersal in terms of reduced desiccation resistance. This trend was preserved in dispersal-selected and non-selected controls as well, where the desiccation cost of dispersal in females was very low compared to the males. Finally, we found that the females instead paid a significant reproductive cost of dispersal.
5. Our results highlight the complex relationship between desiccation stress and dispersal, whereby desiccation resistance can show both a positive and a negative association with dispersal. Furthermore, the sex differences observed in these trait associations may translate into differences in movement patterns, thereby giving rise to sex-biased dispersal.

## 1. Introduction

Biological dispersal is often driven by numerous biotic and abiotic causes that promote movement across space (Matthysen 2012). However, the very process of movement can be costly to the dispersing organisms in several ways (Bonte *et al*. 2012). Investigating the causes and costs of dispersal can therefore help understand the constraints faced by individual organisms (Ronce & Clobert 2012), as well as their potential effects on the population- and community-level consequences of dispersal (Bowler & Benton 2005).

Since dispersal is a key life-history trait in individuals (Bonte & Dahirel 2017), one possible way to decipher its causes and costs is by studying its associations with other life-history and behavioural traits. Collectively known as a ‘dispersal syndrome’ (Ronce & Clobert 2012), these dispersal-trait associations have been documented in several taxa (Stevens *et al*. 2014; Legrand *et al*. 2016; Comte & Olden 2018; Tung *et al*. 2018a). While these trait correlations can help us understand the underlying physiological mechanisms and constraints of dispersal, they are often contingent on the study environment and population history. This is because trait associations change rapidly and significantly if the environment changes, or if the population undergoes evolutionary changes (Chippindale, Ngo & Rose 2003; Jessup & Bohannan 2008; Mishra *et al*. 2018a). Moreover, dispersal may be modulated by many causes at once (Matthysen 2012; Legrand *et al*. 2015), and incur several simultaneous costs to the individuals (Roff 1977; Gros, Hovestadt & Poethke 2008; Bonte *et al*. 2012). Taken together, this makes a thorough investigation of dispersal-trait associations difficult under natural conditions. Therefore, one possibility is to study populations with a known history under a simplified environment to understand how a particular trait association (and hence, the dispersal syndrome) is shaped.

Desiccation stress is one of the factors that can greatly influence dispersal. Not only is it one of the most commonly faced environmental stress for numerous taxa (Black & Pritchard 2002; Holmstrup, Hedlund & Boriss 2002; Kranner *et al*. 2008; Holzinger & Karsten 2013), it is also one of the first signs of an unfavourable environment, as the stress due to lack of water sometimes precedes lack of other resources such as food (Karan & Parkash 1998; Hoffmann & Harshman 1999). Understandably, desiccation not only affects the physiology of individual organisms (e.g. Gibbs, Chippindale & Rose 1997; Folk & Bradley 2004; Bazinet *et al*. 2010), but is also an important determinant of species distributions (e.g. Kellermann *et al*. 2009; Rajpurohit, Nedved & Gibbs 2013). Furthermore, organisms’ responses to desiccation stress are particularly important in the context of climate change and its biological implications (Hoffmann *et al*. 2003; Tuba, Slack & Stark 2011; Van Heerwaarden & Sgrò 2014). Given that dispersal often serves as the first line of defence against unfavourable environments for many taxa (Gerber & Kokko 2018; Riotte-Lambert & Matthiopoulos 2020), it is crucial to investigate the relationship between biological dispersal and desiccation stress.

Desiccation stress can potentially act as both a cause and a cost of dispersal. A high desiccation stress may drive individuals away from an area, while at the same time, the very process of movement can incur desiccation stress to the dispersers. Since males and females in sexually dimorphic species often differ in the amount of body resources and their partitioning along the survival-reproduction axis (Rantala & Roff 2007; Wilkin & Sheldon 2009; Maklakov & Lummaa 2013), differences in their desiccation profiles are commonplace (Jill & Daniel 2003; Matzkin, Watts & Markow 2007; Lyons *et al*. 2014). Similarly, many species exhibit ‘sex-biased dispersal’, a possible reflection of asymmetric cost-benefit outcomes of dispersal between the sexes (Trochet *et al*. 2016; Li & Kokko 2019). While the relationship among environmental stress, dispersal and sex have been recently discussed (Gerber & Kokko 2018), sex differences in the dispersal-desiccation relationship have typically not been studied. This is not surprising given that, investigations into sex differences in dispersal syndromes are relatively rare in the dispersal literature (but see Legrand *et al*. 2016; Mishra *et al*. 2018a). The presence of pervasive sex differences in the life-history and behaviour literature leads us to anticipate some sex differences in the relationship between dispersal and desiccation stress as well. Especially in terms of dispersal costs, it would be interesting to see how the desiccation stress incurred during movement compares with other dispersal-related fitness costs such as female fecundity (Roff & Fairbairn 2007; Guerra 2011).

Here, we investigate the relationship between desiccation stress and dispersal, as well as the associated sex differences, using populations of the common fruit fly (*Drosophila melanogaster*) under controlled environmental conditions. Interestingly, both a positive and a negative association of desiccation stress with dispersal has already been reported in *D. melanogaster* (Mishra *et al*. 2018a), thus making it a suitable system to delineate how the desiccation-dispersal relationship is shaped. Specifically, we asked the following questions: (1) Does desiccation stress act as a cause of dispersal in males and females? (2) Is desiccation stress a cost of dispersal in males and females? (3) Does dispersal evolution alter the desiccation cost of dispersal in either sex, and (4) Do females experience a fecundity cost of dispersal? Our results showed that desiccation stress acts as a significant cause for dispersal for both sexes. However, desiccation stress emerged as a cost of dispersal largely in the males, and was not altered by dispersal evolution. Finally, while the females paid a negligible desiccation cost of dispersal, they experienced a significant cost of dispersal in terms of their fecundity. We discuss these results in the context of *Drosophila* physiology, along with their implications for dispersal patterns.

## 2. Methods

### 2.1 Fly populations

We used large, outbred laboratory populations (breeding size ∼2400 individuals) of *D. melanogaster* for all the experiments in this study. The ancestry of these populations can be traced back to the IV lines, which were wild-caught in South Amherst, MA, USA (Ives 1970). The single-generation experiments in this study were conducted using a baseline population named DB_4_ (Sah, Salve & Dey 2013; Mishra *et al*. 2020). In addition, we used four dispersal-selected populations (namely, VB_1-4_) and their corresponding controls, the non-selected populations (VBC_1-4_), for one experiment. Due to the ongoing selection for higher dispersal every generation, the VB populations have evolved a higher dispersal propensity and ability (Tung *et al*. 2018b), as well as lower desiccation resistance (Mishra *et al*. 2018a), compared with the VBC populations. All the populations were maintained in discrete-generation cycles under uniform environmental conditions of 25 °C temperature and 24-h light.

### 2.2 Dispersal setup

Following previous studies (Mishra *et al*. 2018a; Tung *et al*. 2018b), we used a two-patch dispersal setup for observing fly dispersal. Each dispersal setup comprised a *source* container, a *path* tube and a *destination* container (Fig. 1). In this setup, all the flies for a given treatment/group are first introduced into the *source* container, which opens into a transparent plastic tube (internal diameter ∼1 cm) that serves as the *path*. The other end of the path tube leads into the *destination* container, thereby allowing the dispersal of flies from the *source* to the *destination* container through the *path*, for a fixed duration. Depending on the experiment, the size of the *source* and *destination* containers, as well as the length of the *path* tube, can be customized. A single experiment typically involves multiple such dispersal setups, maintained under uniform environmental conditions. At the end of a dispersal run, these dispersal setups are dismantled, and the flies found in each part (*source/path/destination*) are used as per the experimental requirements.

**Fig. 1:**
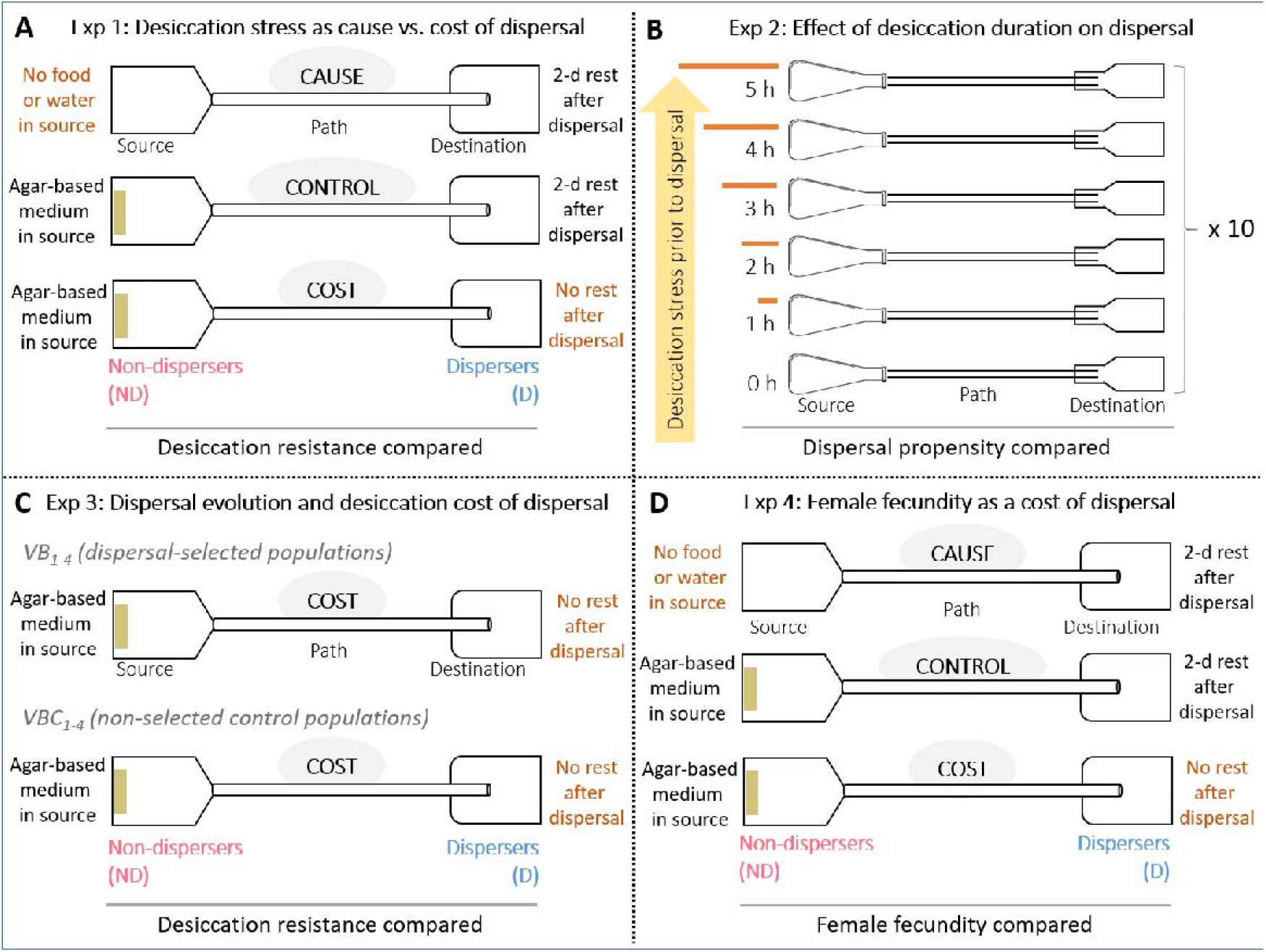
Schematics of the experimental design. (A) Experiment 1 investigated the role of desiccation stress as a cause vs. cost of dispersal. Using a *source-path-destination* setup, age-matched flies from an outbred baseline population (DB_4_) were segregated into non-dispersers (ND) and dispersers (D) under three scenarios: *Cause* (no food or water in *source*; rest provided after dispersal run), *Control* (agar-based banana-jaggery medium in *source*; rest provided after dispersal run), and *Cost* (agar-based banana-jaggery medium in source; no rest provided after dispersal run). ND and D flies within each scenario were then assayed for their desiccation resistance. (B) Experiment 2 further examined the role of desiccation stress as a cause of dispersal. Groups of age-matched flies from DB_4_ population were subjected to different durations of desiccation stress (0–5 h) before being subjected to dispersal assay. (C) Experiment 3 investigated whether the desiccation *Cost* of dispersal differs between populations selected for higher dispersal (VB_1–4_) and their non-selected controls (VBC_1–4_). Desiccation resistance of all eight population blocks was compared under the *Cost* scenario similar to Experiment 1. (D) Experiment 4 examined the role of female fecundity as a cause vs. *Cost* of dispersal. Here, female ND and D flies for the three scenarios (*Cause, Control*, and *Cost*) were assayed for their fecundity.

### 2.3 Experiments

We carried out a series of experiments to address various questions related to Causes and Costs of dispersal. The protocols, type of data obtained and the statistical analyses are presented separately for each experiment below.

#### 2.3.1 Experiment 1: Desiccation stress as Cause vs. *Cost* of dispersal

We first examined whether desiccation stress acts as a Cause and emerges as a Cost of dispersal in *D. melanogaster*. For this, we started with ∼19,200 age-matched (12-day-old from egg collection) adult flies from the DB_4_ population that were reared under identical conditions of *ad libitum* food and water. Cylindrical, translucent plastic containers (∼1.5 L volume) were used as *source* and *destination*, along with a *path* length of 6 m, to assemble two-patch dispersal setups (described in section 2.2). Batches of the aforementioned DB_4_ individuals were then introduced into eight such dispersal setups (∼2400 individuals per setup) and allowed to disperse for 5 h. By modulating two factors, i.e. presence of agar-based food (banana-jaggery medium) in the *source* container, and the provision of rest to flies after the dispersal run, we devised three scenarios (Fig. 1A, see explanation in next paragraph): (a) *Cause* scenario, where we could identify whether desiccation stress was a Cause of dispersal, (b) *Control* scenario, where desiccation stress was expected to be neither a Cause nor a Cost of dispersal, and (c) *Cost* scenario, where we could identify whether desiccation stress was a *Cost* of dispersal (Fig. 1A). In each of the three scenarios, the flies that completed dispersal from the *source* to the *destination* were termed as dispersers (D), whereas the flies that were found inside the *source* container were termed as non-dispersers (ND). The flies found in the path at the end of the dispersal run were not used in this experiment.

In the *Cause* scenario, there was no food or water in the *source*, making desiccation stress a likely driver of dispersal away from the source. After the dispersal event, we collected the ND and D flies separately and provided them a 2-day rest with *ad libitum* food and water, so that the D flies could recuperate any energy Costs of dispersal run. Thereafter, we assayed 200 ND and 200 D flies (100 males+100 females each) for their desiccation resistance (Supplementary Text S1.1), to assess whether they differed in terms of their inherent desiccation sensitivity (Fig. 1A: *Cause* scenario). Here the assumption is that the rest of 2 days is sufficient to ameliorate any negative effects on desiccation sensitivity (similar to Mishra *et al*. 2018a).

In the *Control* scenario, we provided agar-based banana-jaggery medium in the *source* container during the dispersal run, thereby removing desiccation stress as a possible driver of dispersal. Similar to the *Cause* scenario, the dispersal event was followed by a 2-day rest to both ND and D flies, to offset any energy Costs of dispersal (Fig. 1A: *Control* scenario). Subsequently, we compared the desiccation resistance of 200 ND and 200 D flies, to ascertain if there were any unaccounted-for differences between them, i.e. other than those detected in *Cause* and *Cost* scenarios.

The *Cost* scenario was complementary to the *Cause* scenario. Here, we provided banana-jaggery medium in the *source* container, thereby removing desiccation stress as a Cause of dispersal, but did not allow any rest after dispersal. As above, we then compared the desiccation resistance of 200 ND and 200 D flies, with any difference attributed to the energy Costs of dispersal (Fig. 1A: *Cost* scenario).

The statistical analyses for this experiment, as well as those described in subsequent sections, were carried out in R v4.0.3 (R Core Team 2020). Here, the desiccation data from Experiment 1 were analysed together in a single mixed-model GLM using the ‘lmer’ function from the ‘lme4’ package v1.1-25 (Bates *et al*. 2015), with *scenario* (*Cause*/*Control*/*Cost*), *dispersal* (ND/D) and *sex* (male/female) as the fixed factors. As the flies were assayed in single-sex groups of 10 individuals within a vial (Supplementary Text S1.1), we included *vial identity* (1–10) as a random factor that was nested within the *scenario × dispersal × sex* interaction. Following a Type III analysis of deviance to ascertain the significance of the fixed factors and their interactions in GLM via the ‘Anova’ function in ‘car’ package v3.0-10 (Fox & Weisberg 2019), we carried out the relevant pairwise comparisons using the ‘pairs’ function in the ‘emmeans’ package v1.5.2-1 (Lenth 2020). Cohen’s d was used as a measure of effect size for significantly different pairs of means, with the effect interpreted as large, medium, and small for d ≥ 0.8, 0.8 > d ≥ 0.5, and d < 0.5, respectively (Cohen 1988).

#### 2.3.2 Experiment 2: Effect of desiccation duration on dispersal

Here, we investigated how dispersal changes with the duration of desiccation stress. For this, we segregated age-matched (12-day-old from egg collection) DB_4_ flies into multiple groups of 120 individuals (60 males + 60 females) that were subjected to varying durations of desiccation stress (0, 1, 2, 3, 4, and 5 h) before being subjected to dispersal assay in separate dispersal setups (Fig. 1B). The *source* here was a 100-mL glass flask without any food or water, the *path* length was 2 m, and the destination was a 250-mL plastic bottle. The dispersal assay lasted for 2 h. Following a previous protocol (Mishra *et al*. 2018b; Mishra, Chakraborty & Dey 2020; Mishra *et al*. 2020), the experiment was carried out over 10 consecutive days with a fresh set of age-matched flies every day. This allowed us to assay one replicate of every desiccation treatment each day, yielding 10 replicates blocked by *day*. In total, 6000 flies (5 desiccation treatments × 2 sexes × 10 days × 60 flies treatment^-1^ sex^-1^ day^-1^) were assayed for this experiment. From the dispersal assay, we collected data on dispersal propensity (i.e. the fraction of flies that dispersed from the source: Friedenberg 2003; Supplementary Text S1.2). To account for any day-to-day micro-environmental variation, we used *day* as a random blocking factor in the analysis. Therefore, the dispersal propensity data were analysed in a mixed model binomial GLM (with logit link function) using the ‘glmer’ function from the ‘lme4’ package v1.1-25 (Bates *et al*. 2015), with *desiccation duration* (0, 1, 2, 3, 4, and 5 h) and *sex* (male and female) as fixed factors, and *day* (1–10) as the random factor. Following analysis of deviance via ‘Anova’ function in ‘car’ package v3.0-10 (Fox & Weisberg 2019), appropriate pairwise comparisons were carried out using the ‘pairs’ function in ‘emmeans’ package v1.5.2-1 (Lenth 2020).

#### 2.3.3 Experiment 3: Dispersal evolution and desiccation Cost of dispersal

Here, we used dispersal-selected populations (VB_1-4_), which have a higher dispersal propensity and travel longer distances (Tung *et al*. 2018b), as well as a lower desiccation resistance, than their non-selected Controls (VBC_1-4_) (Mishra *et al*. 2018a). In this experiment, we investigated whether the VB and VBC populations differ in their desiccation Cost of dispersal. This would help determine if selection for dispersal under desiccated conditions has altered the magnitude of proximate Cost paid by dispersers. We subjected ∼2400 age-matched individuals per population block (1–4) of each population type (VB/VBC) to segregation into ND and D individuals under the Cost scenario as described in section 2.3.1 (Fig. 1C). Thereafter, we assayed 100 males and 100 females (in groups of 10 individuals/vial) from each of the eight populations (VB_1-4_ and VBC_1-4_) for their desiccation resistance (Supplementary Text S1.1). The entire desiccation resistance data were analysed using a mixed-model GLM with the ‘lmer’ function in ‘lme4’ package v1.1-25 (Bates *et al*. 2015), with *dispersal selection* (VB/VBC), *dispersal* (ND/D) and *sex* (male/female) as fixed factors, and *population block* (1–4) and *vial identity* (1–10) as random factors. Here, *vial identity* was nested inside the *dispersal selection × dispersal × sex × population block* term. Following the GLM, we used the ‘Anova’ function in ‘car’ package v3.0-10 (Fox & Weisberg 2019) for analysis of deviance, and subsequently, the ‘pairs’ function in ‘emmeans’ package v1.5.2-1 (Lenth 2020) for relevant pairwise comparisons.

#### 2.3.4 Experiment 4: Female fecundity as Cause vs. Cost of dispersal

This experiment aimed to examine whether females paid a dispersal Cost in terms of their fecundity. The female flies in this experiment were from the same ND and D groups of flies that were segregated in Experiment 1, giving rise to: (a) *Cause* scenario, defined by the lack of suitable oviposition site in *source* container, (b) *Control* scenario, with suitable oviposition surface (i.e. banana-jaggery medium) in the *source* and provision of rest after dispersal run, and (c) *Cost* scenario, where no rest is provided and flies were assayed for their fecundity immediately after dispersal (Fig. 1D). We counted the female fecundity as the number of eggs laid over a 12-h period, with the ND and D flies for each scenario assayed together (see Supplementary Text S1.3 for details). The entire fecundity data were analysed together with a quasi-Poisson GLM (with log link function) using the ‘glm’ function in ‘stats’ package v4.0.3 (R Core Team 2020), with *scenario* (*Cause, Control*, and *Cost*) and *dispersal* (ND and D) as the fixed factors. As above, we used the ‘Anova’ function in ‘car’ package v3.0-10 (Fox & Weisberg 2019) for analysis of deviance, and the ‘pairs’ function in ‘emmeans’ package v1.5.2-1 (Lenth 2020) for relevant pairwise comparisons.

## 3. Results

### 3.1 Desiccation stress as a Cause vs. Cost of dispersal

Desiccation resistance data from Experiment 1 showed a significant *scenario × dispersal × sex* interaction (χ^2^ = 7.20, p = 0.027). Analysis of pairwise differences for this interaction revealed a number of results (Supplementary Text S2.1). First, there was no difference in the desiccation resistance of dispersers vs. non-dispersers in the *Control* scenario (p_males_ = 0.16; p_females_ = 0.34) (Fig. 2B, 2E). This was expected, as all these flies had access to *ad libitum* food and water in the source container, as well as a 2-day rest after the dispersal event. Second, dispersers in the Cause scenario had a lower desiccation resistance than non-dispersers (p_males_ = 0.006, d = 1.63 (large); p_females_ = 0.005, d = 1.10 (large)) (Fig. 2A, 2D). This implies that desiccation stress likely served as a Cause of dispersal in both sexes. Third, while males experienced a Cost of dispersal in terms of their desiccation resistance (p < 10^−4^, d = 1.21 (large)), no such Cost was seen in females (p = 0.86) (Fig. 2C, 2F).

**Fig. 2:**
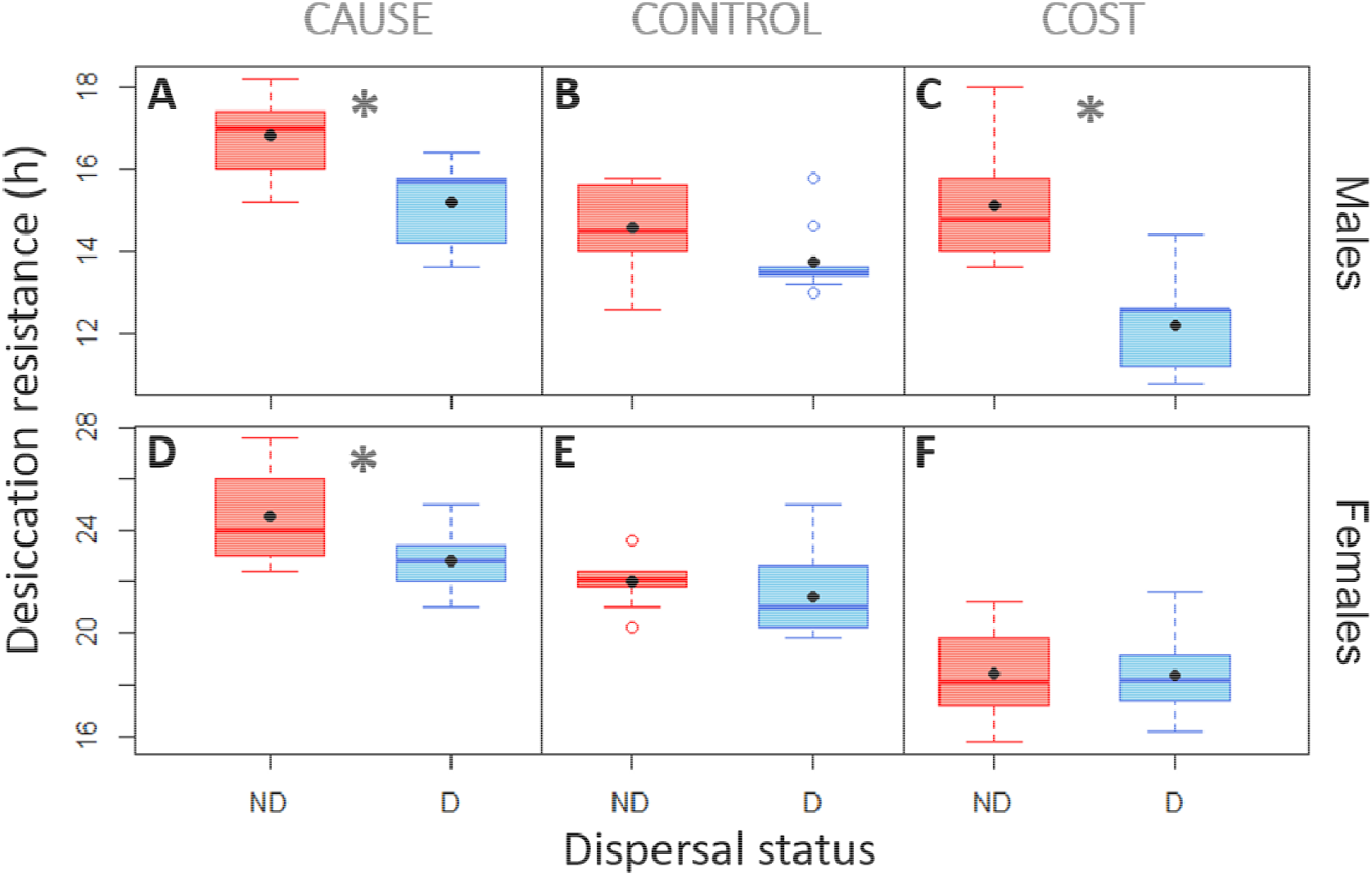
Desiccation stress as Cause vs. Cost of dispersal (Experiment 1). Desiccation resistance for non-disperser (ND) and disperser (D) flies from an outbred, baseline population (DB_4_), under three scenarios: *Cause, Control*, and *Cost*. Data for males and females are presented in the top and bottom rows, respectively. Edges of the boxplots represent 25^th^ and 75^th^ percentiles of the data. The black dots represent means and the lines inside box represent medians. Asterisks (*) indicate a significant difference (p < 0.05) between ND and D flies within a given panel. Note that the scales on Y-axis differ between the males and the females. See Supplementary Text S2.1 for the exact p values.

### 3.2 Desiccation stress as a Cause of dispersal in both sexes

The role of desiccation stress as a Cause of dispersal was further investigated in Experiment 2. Analysis of data from this experiment revealed that the *desiccation duration × sex* interaction was significant (χ^2^ = 17.10, p = 0.004), indicating an asymmetric effect of desiccation duration on dispersal propensity of males and females. However, pairwise comparisons revealed an increasing trend of dispersal propensity with longer durations of desiccation stress in both sexes, with a somewhat greater effect observed in males (Fig. 3) (Supplementary Text S2.2). Therefore, the results from both Experiments 1 and 2 suggested that desiccation stress served as a Cause of dispersal in both sexes, with longer durations of desiccation leading to greater dispersal.

**Fig. 3:**
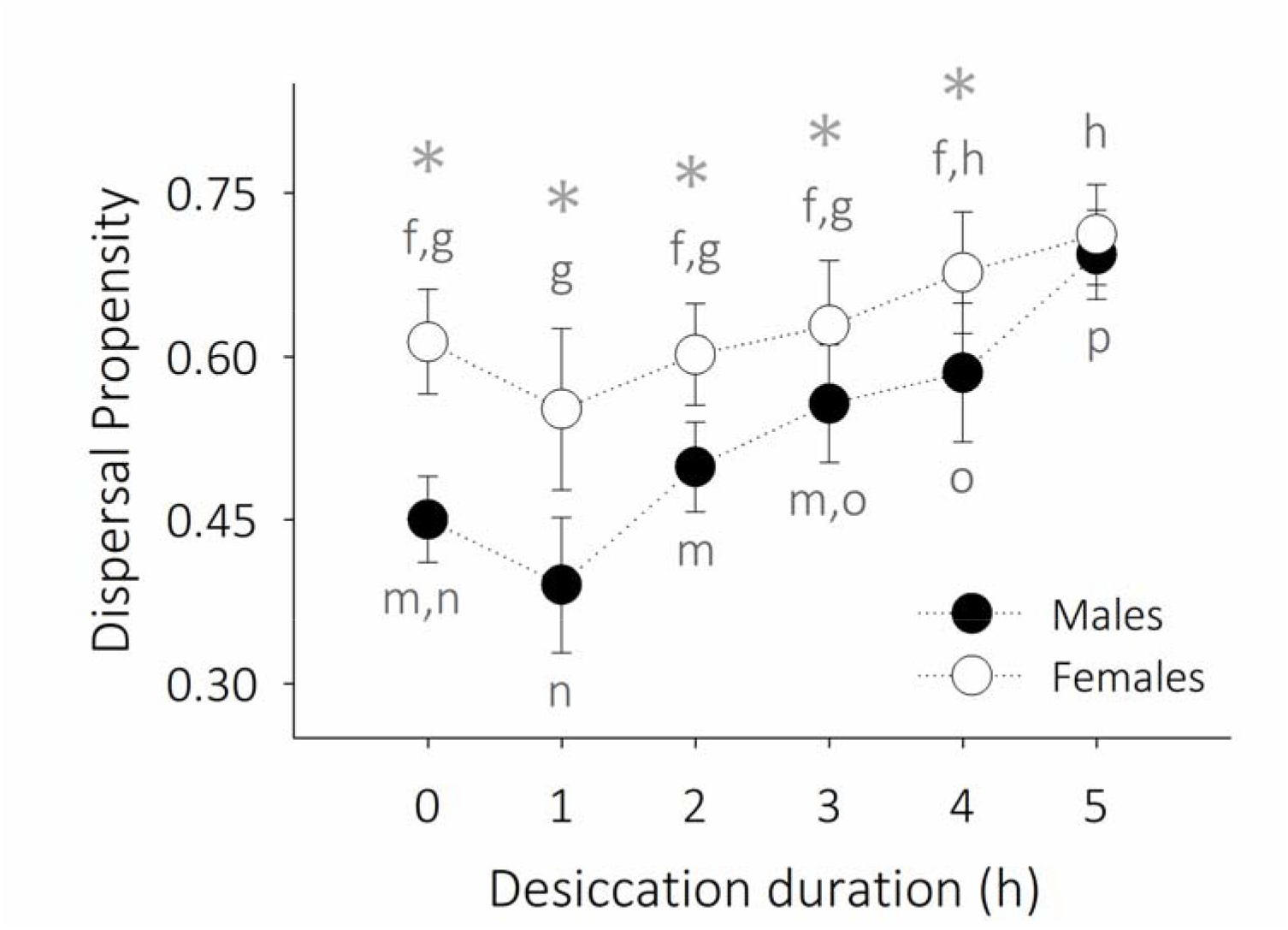
Effect of desiccation duration on dispersal propensity (Experiment 2). Dispersal propensity (± SE) for age-matched flies from an outbred baseline population (DB_4_) subjected to desiccation stress for different durations (0–5 h). Each point represents the average of 10 replicates (each with 120 individuals). For a given sex, the changes in dispersal are examined by comparing the propensity means across the six desiccation durations (significant differences denoted using different lower-case letters: starting with m for males and f for females). Asterisks (*) denote a significant difference in male and female dispersal for a given desiccation duration. See Supplementary Text S2.2 for the exact p values.

### 3.3 Desiccation stress as a sex-biased Cost of dispersal

Next, we examined the role of desiccation stress as a Cost of dispersal using four dispersal-selected populations (VB_1-4_) and their corresponding non-selected Controls (VBC_1-4_) (Experiment 3). Desiccation resistance data from this experiment revealed a significant *dispersal × sex* interaction (χ^2^ = 9.52, p = 0.002), with males experiencing a relatively larger desiccation *Cost* of dispersal (p < 10^−4^, d = 1.86 (large)) (Fig. 4A, 4B) than females (p < 10^−4^, d = 0.42 (small)) (Fig. 4C, 4D). Moreover, the *dispersal selection × dispersal* (χ^2^ = 2.33, p = 0.13) and *dispersal selection × dispersal × sex* (χ^2^ = 0.19, p = 0.66) interactions were not significant, indicating that this result was consistent for both Control (VBC) and dispersal-selected (VB) populations (Supplementary Text S2.3).

**Fig. 4:**
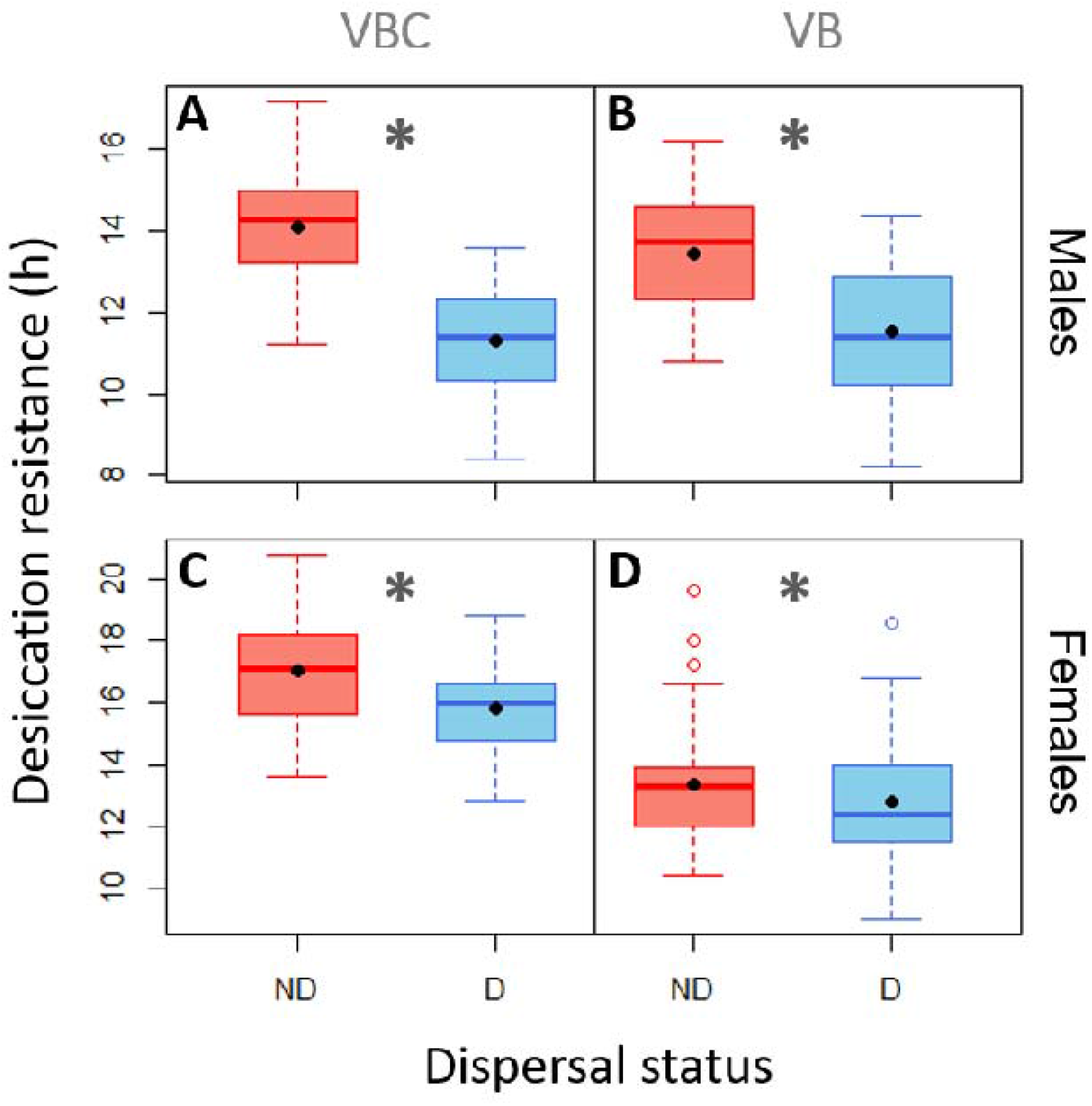
Dispersal evolution and desiccation Cost of dispersal (Experiment 3). Desiccation resistance of non-dispersers (ND) and dispersers (D) from VB_1-4_ (dispersal-selected) and VBC_1-4_ (*Control*) populations. Data for males and females are presented in the top and bottom rows, respectively. Edges of the boxplots represent 25^th^ and 75^th^ percentiles of the data. Asterisks (*) indicate a significant difference (p < 0.05) between ND and D flies within a given panel. See Supplementary Text S2.3 for the exact p values.

### 3.4 Significant Cost of dispersal for females in terms of fecundity

As minimal or no desiccation Cost of dispersal was observed for females (Sections 3.1 and 3.3), we investigated if there was a reproductive Cost of dispersal for the females (Experiment 4). Analysis of the female fecundity data from this experiment (presented in Supplementary Text S2.4) revealed a significant *scenario × dispersal* interaction (χ^2^ = 17.90, p = 0.0001). Pairwise comparisons for this interaction revealed no significant difference between dispersers and non-dispersers under the *Control* (p = 0.63) (Fig. 5B) and Cause (p = 0.16) scenarios (Fig. 5A), but a significant difference in the *Cost* scenario: disperser females had a lower fecundity than non-disperser females (p = 0.0001, d = 0.68 (medium)) (Fig. 5C). Therefore, we concluded that female flies pay a *Cost* of dispersal in terms of their fecundity.

**Fig. 5:**
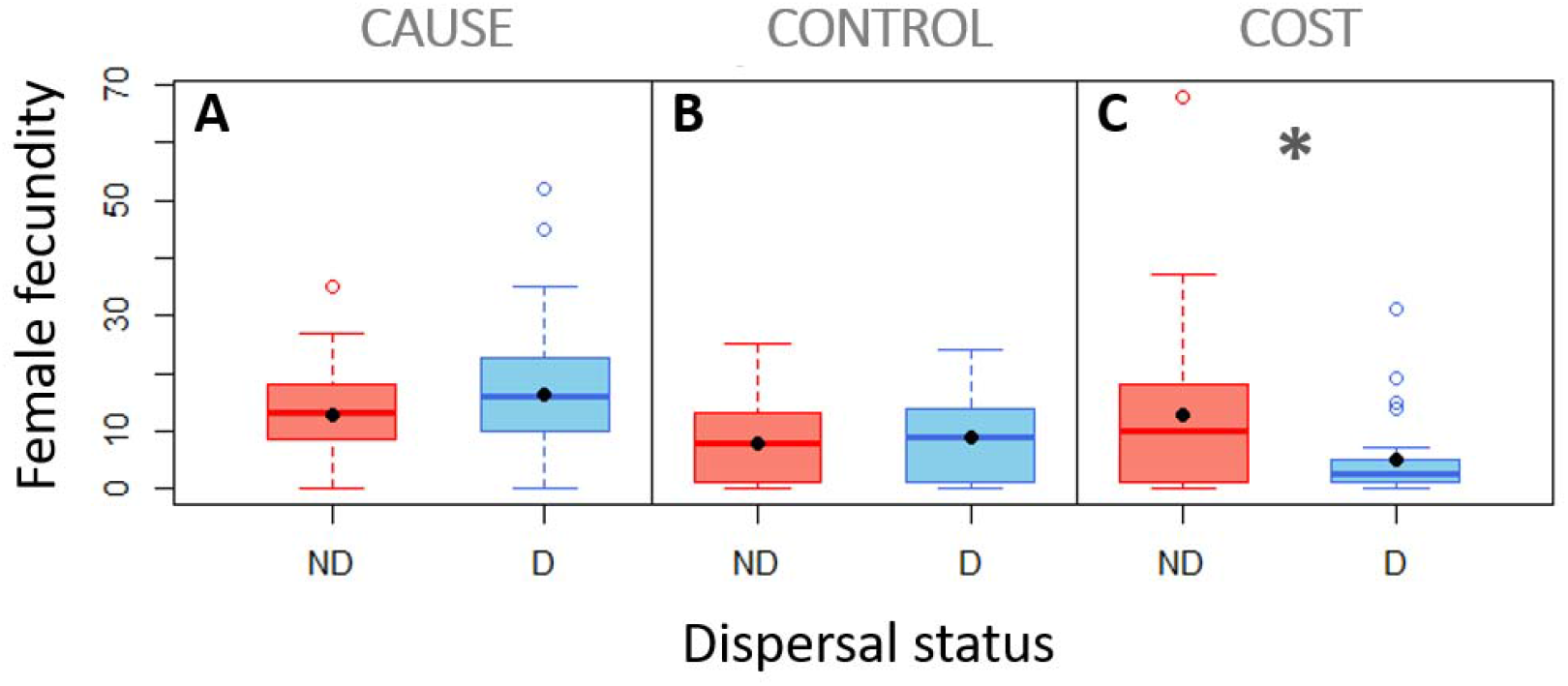
Female fecundity as Cause vs. Cost of dispersal (Experiment 4). Female fecundity for non-disperser (ND) and disperser (D) flies from an outbred, baseline population (DB_4_), under three scenarios: *Cause, Control*, and *Cost*. Edges of the boxplots represent 25^th^ and 75^th^ percentiles of the data. Asterisks (*) indicate a significant difference (p < 0.05) between ND and D flies within a given panel. See Supplementary Text S2.4 for the exact p values.

## 4. Discussion

### 4.1 Desiccation stress as a Cause of dispersal in both sexes

Environmental stress, among other things, can serve as a major Cause of biological dispersal. At the same time, the very process of dispersal can be stressful to individuals. When monitored after a dispersal event, the stress-resistance ability of organisms are often found to be lower (Graves *et al*. 1992). This decrease can come about in three different ways. First, the dispersers might be the ones that were more susceptible to the stress, and hence they dispersed. Second, even if the stress resistance of the dispersers is inherently similar to that of the non-dispersers, the energy spent in the act of dispersal reduces the stress-resistance ability of the former. Third, it might be an interaction of the above two scenarios. Unfortunately, these questions are very difficult to answer, particularly when there is no *a priori* way of distinguishing between a disperser and a non-disperser. Here, we investigated this complex relationship, using desiccation as the type of stress and fruit flies as a model system. Our experimental design allowed us to explicitly *Control* for other confounds when a particular aspect of the desiccation-dispersal relationship was being examined.

To begin with, Experiment 1 revealed that the disperser (D) flies had a lower desiccation resistance than the non-disperser (ND) flies under the *Cause* scenario (Figs. 2A and 2D). Comparing the results with the *Control* scenario, which showed no difference between ND and D flies (Figs. 2B and 2E), we could conclude that desiccation stress indeed served as a significant driver of dispersal for both male and female flies. This is in line with the expectation from literature that dispersal is one of the foremost ways for escaping unfavourable conditions (Gerber & Kokko 2018), not only in animal taxa (Cremer & Heinze 2003; Riotte-Lambert & Matthiopoulos 2020) but also in plants (Martorell & MartínezCLópez 2014). While this is not a surprising result, our study demonstrates it explicitly using a unique setup, where we were able to Control for the possible confound of desiccation as a Cost of dispersal (Fig. 1A).

Going a step further, we demonstrate in Experiment 2 how *Drosophila* dispersal changes with increasing desiccation stress (Fig. 3). Given that desiccation resistance is highly correlated with glycogen content in fruit flies (Gibbs, Chippindale & Rose 1997), one might have expected a decrease in dispersal at longer desiccation durations, where the flies likely faced a severe depletion of their glycogen reserves (Folk & Bradley 2004; Bazinet *et al*. 2010). Surprisingly however, this was not the case in Experiment 2, where flies of both sexes showed a nearly monotonic increase in their dispersal propensity with increasing desiccation stress (Fig 3). This means that, at least for the duration of desiccation stress (up to 5 h) imposed in Experiment 2, the flies were in a state to successfully initiate dispersal. However, as a corollary, it also means that organisms likely do not disperse until the stress turns acute, which may make them more susceptible to dispersal-related risks and Costs (see Section 4.2). It is possible that this delay in emigration could be a function of how long it takes to initiate the stress physiological response. Overall, we speculate that the ability to perceive stress would play a role in shaping the dispersal-mediated escape response from stressful habitats.

Since dispersal is also known to incur various Costs (reviewed in Bonte *et al*. 2012), the process of dispersal itself can induce stress or increase the susceptibility of dispersing individuals to stress. We explored the potential desiccation Cost of dispersal using the *Cost* scenario, in Experiments 1 and 3.

### 4.2 Sex-biased *Cost* of dispersal in terms of desiccation stress

Given that active dispersal involves expenditure of energy, it is likely that flies spend a part of their glycogen reserves during dispersal (Graves *et al*. 1992), which can reduce their desiccation resistance following a dispersal event. Experiment 1 confirmed a Cost of dispersal in terms of their desiccation resistance, although it was not symmetric between the two sexes. A significant desiccation Cost of dispersal was observed for males (Fig. 2C) but not for females (Fig. 2F) in the DB_4_ population. Similarly, Experiment 3 revealed that the desiccation *Cost* of dispersal was much higher in males (Fig. 4A, 4B) than in females (Fig. 4C, 4D) (see section 3.3 for the exact effect sizes). As both dispersal-selected (VB) and non-selected Control (VBC) flies showed a male-biased desiccation Cost, we concluded that the evolution of dispersal did not alter the immediate desiccation Cost of dispersal between these populations.

A potential explanation for the sex bias in desiccation Cost is the sexual dimorphism in body size and desiccation resistance of *D. melanogaster* adults. A positive association between desiccation resistance and body size is well documented in adult fruit flies (Parsons 1970; Clark & Doane 1983). Given that female fruit flies are typically larger than their male counterparts, they typically tend to have a higher desiccation resistance as well (Gibbs, Chippindale & Rose 1997; Matzkin, Watts & Markow 2007; Mishra *et al*. 2018a). As a result, the females likely had greater resources to begin with, which allowed them to successfully undertake dispersal without paying a high desiccation *Cost*. This is also congruent with the observation that dispersal evolution has not led to a change in the body size of VB females relative to their VBC Controls (Mishra *et al*. 2018a; Tung *et al*. 2018a).

It is possible that the dispersal Cost for females manifests not in terms of their somatic maintenance (here, desiccation resistance), but instead their reproductive potential. This is in line with the results of several life-history studies on trade-offs that show a reproductive Cost instead of somatic Costs in females (Miyatake 1997; Ghalambor & Martin 2001; Djawdan *et al*. 2004; Muller-Landau 2010). In such cases, female fecundity is often one of the first traits to exhibit this Cost. Given the energy-intensive nature of active dispersal (as evidenced by the dispersal Cost borne by males in this study), female fecundity could show a Cost of dispersal. Therefore, we next investigated the association between female fecundity and dispersal.

### 4.3 Fecundity Cost of dispersal for female flies

The relationship between dispersal and fecundity varies across taxa. A negative association between dispersal and fecundity has been reported in several wing-dimorphic insects (reviewed in Guerra 2011), wing-monomorphic insects (reviewed in Tigreros & Davidowitz 2019), as well as other taxa such as *C. elegans* (Friedenberg 2003). These results are typically explained as a developmental or energetic *Cost* of dispersal in terms of fecundity. In contrast, a positive association between dispersal and fecundity has been observed in many mammalian taxa (reviewed in Stevens *et al*. 2014). Here, the typical explanation is twofold. First, individuals with better body condition, including higher fecundity, could be better able to complete dispersal. Second, high fecundity could lead to high dispersal via increased kin competition in a given habitat. Of course, it is also possible that the dispersal-fecundity relationship, like other dispersal-trait associations, is modulated by the environmental context (e.g. Legrand *et al*. 2016; Mishra *et al*. 2018a). For instance, the fecundity Cost of dispersal may be particularly strong under limiting resources. Similarly, the positive association between dispersal and fecundity might be altered by the population density and level of resources in the originating patch (e.g. Einum, SundtCHansen & H. Nislow 2006). Therefore, experiments under *Control*led conditions, which can take the ecological context into account, can provide important insights into the relationship between fecundity and dispersal.

Experiment 4 revealed that, while there was no difference under the *Cause* and *Control* scenarios, D females had a significantly lower fecundity than ND flies in the *Cost* scenario (Fig. 5). What makes our result interesting is that females showed a fecundity *Cost* before the somatic *Cost* of dispersal, at least in terms of desiccation resistance (*cf*. Figs. 2F and FC). A plausible explanation for this is that, under stressful conditions, individuals may prioritize survival over potential reproduction. This has been observed in other life-history traits as well, where allocation of resources into somatic maintenance can, at times, take priority over reproductive investment (Djawdan *et al*. 2004; Muller-Landau 2010; Martorell & MartínezCLópez 2014). In particular, given that dispersal is a key life-history trait (Bonte & Dahirel 2017) with several potential Costs (Bonte *et al*. 2012), the fecundity trade-off observed here is in line with the observations for other wing-monomorphic insects (Tigreros & Davidowitz 2019).

### 4.4. Implications

Our results revealed desiccation as a Cause of dispersal for both sexes in *Drosophila melanogaster*, and dispersal propensity of both male and female flies increased with increasing desiccation duration. In addition, we observed a male-biased Cost of dispersal in terms of desiccation resistance, while the female flies paid a fecundity Cost of dispersal. We discuss some implications of our results below.

First, these results demonstrate that the relationship between stress and dispersal is likely complicated. On one hand, stress is likely to drive dispersal of individuals away from an area. On the other hand, dispersing individuals incur a further Cost of dispersal in terms of increased stress. Therefore, early dispersers from a population may be the least stress-tolerant individuals. In contrast, highly stress-tolerant individuals could delay emigration in response to a stress. As a result, if dispersal occurs across habitats with high connectivity, stress-intolerant individuals may have the highest dispersal propensity (e.g. Fig. 3). However, if the inter-habitat connectivity is poor, only the relatively stress-resistant individuals in a population would be able to undertake dispersal successfully by surviving the large dispersal Costs.

Second, sex differences in the somatic Costs of dispersal may effectively lead to instances of sex-biased dispersal, even if a similar number of male and female individuals emigrate from a given area. This is beCause the stress-sensitive sex (e.g. males in the current study) may not be able to complete dispersal as successfully as the stress-resistant sex (here, females). As a result, in the species where mating occurs after a dispersal event, such differences can lead to a skew in the local sex ratio of the dispersed population and consequently mate limitation.

Moreover, the sex-biased nature of dispersal Costs can result in demographic consequences through dispersal syndromes (Mishra *et al*. 2018a; Shaw, Kokko & Neubert 2018). For instance, if the fecundity of immigrant females in a new area is reduced as a consequence of dispersal, then they may not be able to compete with the resident females in that area. As a result, the apparent prioritization of fitness *Cost* over somatic *Cost* in females, as observed here, can hamper their settlement ability in a new habitat.

Finally, while dispersal is often considered an effective escape route against environmental stress (Boeye *et al*. 2013; Travis *et al*. 2013), it might not be enough to offset the fitness reduction *Cause*d by changing climatic conditions (Buckley, Tewksbury & Deutsch 2013). The situation might worsen further with dispersal-associated Costs that hamper the stress-handling ability of individuals and their biological fitness (Cheptou *et al*. 2008).

Consequently, there is a need to incorporate information on the physiological condition of dispersers in models that consider dispersal as a mode of escape from stressful habitats.

## Acknowledgements

We thank Mohammed Aamir Sadiq and Sahana Srivathsa for help with the experiments. AM and ST were supported by a Senior Research Fellowship from the Council of Scientific and Industrial Research, Government of India. VRSS was supported through the GE Foundation Scholar Leaders Program. PMS was supported through the INSPIRE fellowship of Department of Science and Technology (DST), Government of India. This study was supported by a research grant (#CRG/2018/001333) from Science and Engineering Research Board, DST, Government of India, and internal funding from IISER-Pune.

## Authors’ Contributions

AM and SD conceived the ideas and designed methodology; AM, ST, VRSS and PMS collected the data; AM analyzed the data; AM and SD led the writing of the manuscript. All authors contributed critically to the drafts and gave final approval for publication.

## Supporting Information for

## Text S1. Assay details

### Text S1.1 Desiccation resistance assay (Experiments 1 and 3)

Desiccation resistance for a fly was measured as the duration that it could survive without food and moisture. To quantify this, same-sex groups of 10 flies each were introduced into empty transparent vials and monitored until the death of the last fly in each vial, in a well-lit environment maintained at 25 °C. The survivorship checks were conducted every 2 hours, and 10 such replicate vials were used per sex.

### Text S1.2 Dispersal assay (Experiment 2)

For every two-patch dispersal setup (replicate), we counted the number of male and female flies that reached the destination during each of the 15-min intervals until the end of dispersal assay (2 h). In addition, we recorded the number and sex of flies that emigrated from the source but did not reach the destination, i.e. those found within the path tube at the end of the dispersal assay.

These data were used to estimate the dispersal propensity, i.e. the proportion of flies that initiated dispersal from the source, as:

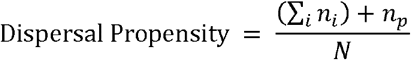

where *n_i_* is the number of flies that reached the destination during the *i*^th^ 15-min interval, *n_p_* is the number of flies found within the path at the end of dispersal assay and *N* is the total number of flies introduced in the setup (here, 120).

### Text S1.3 Fecundity assay (Experiment 4)

Female fecundity was assessed as the number of eggs laid per female over a 12-h period. The flies were anaesthetized under mild CO_2_ and pairs of one male and one female each were introduced into individual 50-mL centrifuge tubes containing a banana-jaggery food cup. The tube had provision for aeration and the food in the food cup provided a surface for laying eggs. Forty such replicates were set up per group (i.e. dispersers/non-dispersers) per scenario. The setups were left undisturbed for 12 hours in a well-lit environment maintained at 25 °C. At the end of 12 hours, the flies were discarded, and the eggs laid on the food were counted under a stereo microscope.

## Text S2. Detailed statistical analyses

### Text S2.1 Experiment 1

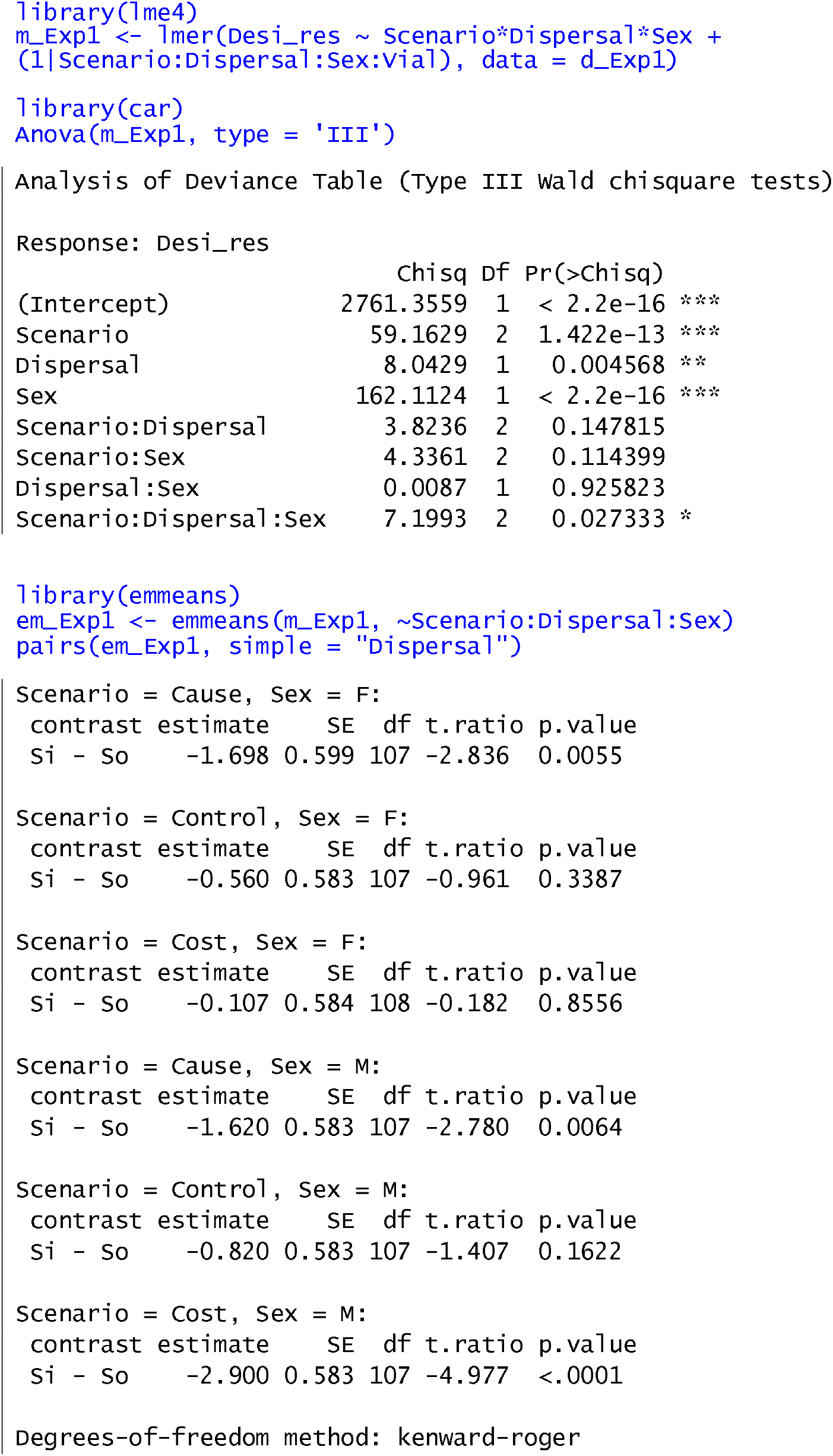

### Text S2.2 Experiment 2

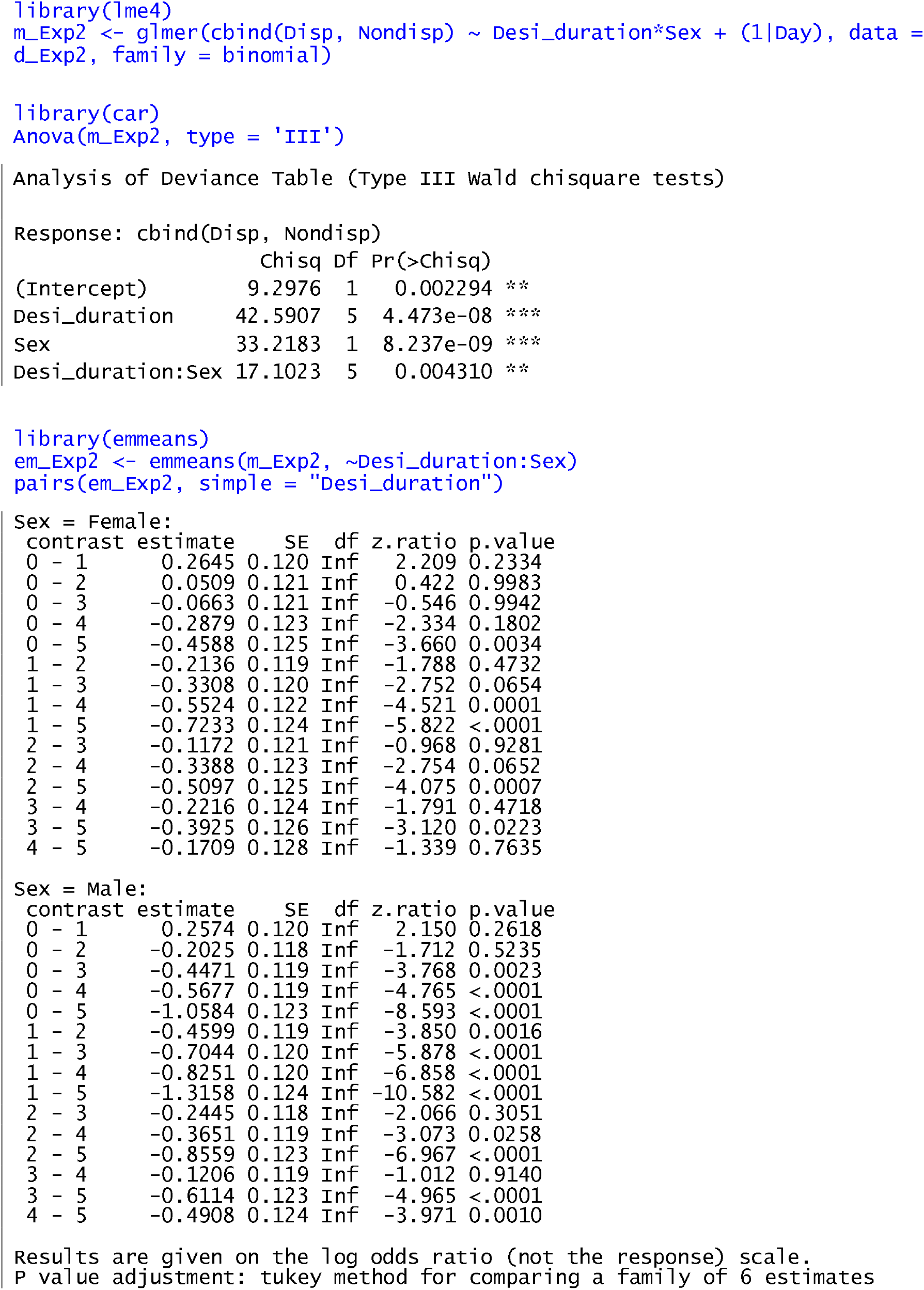
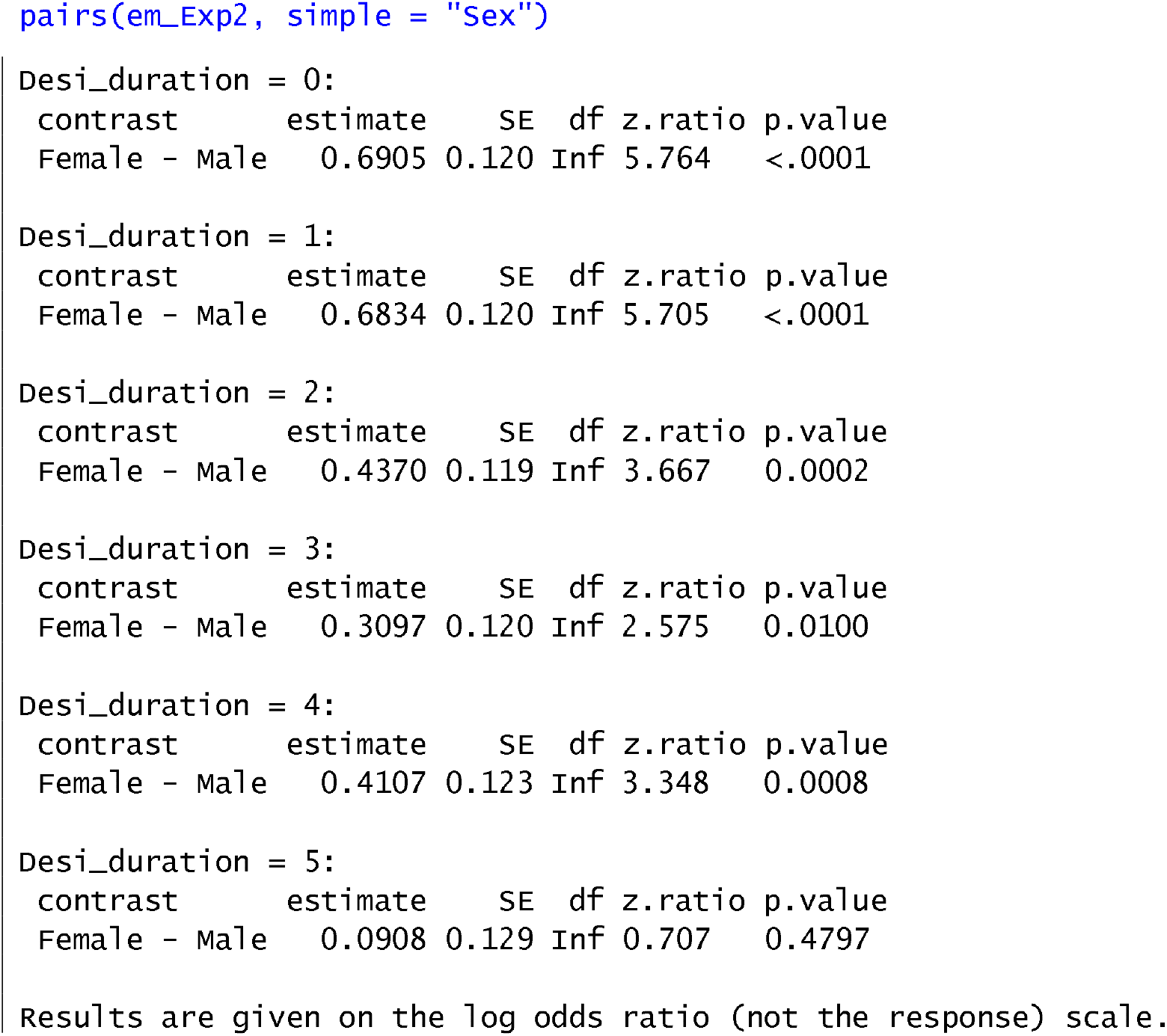

### Text S2.3 Experiment 3

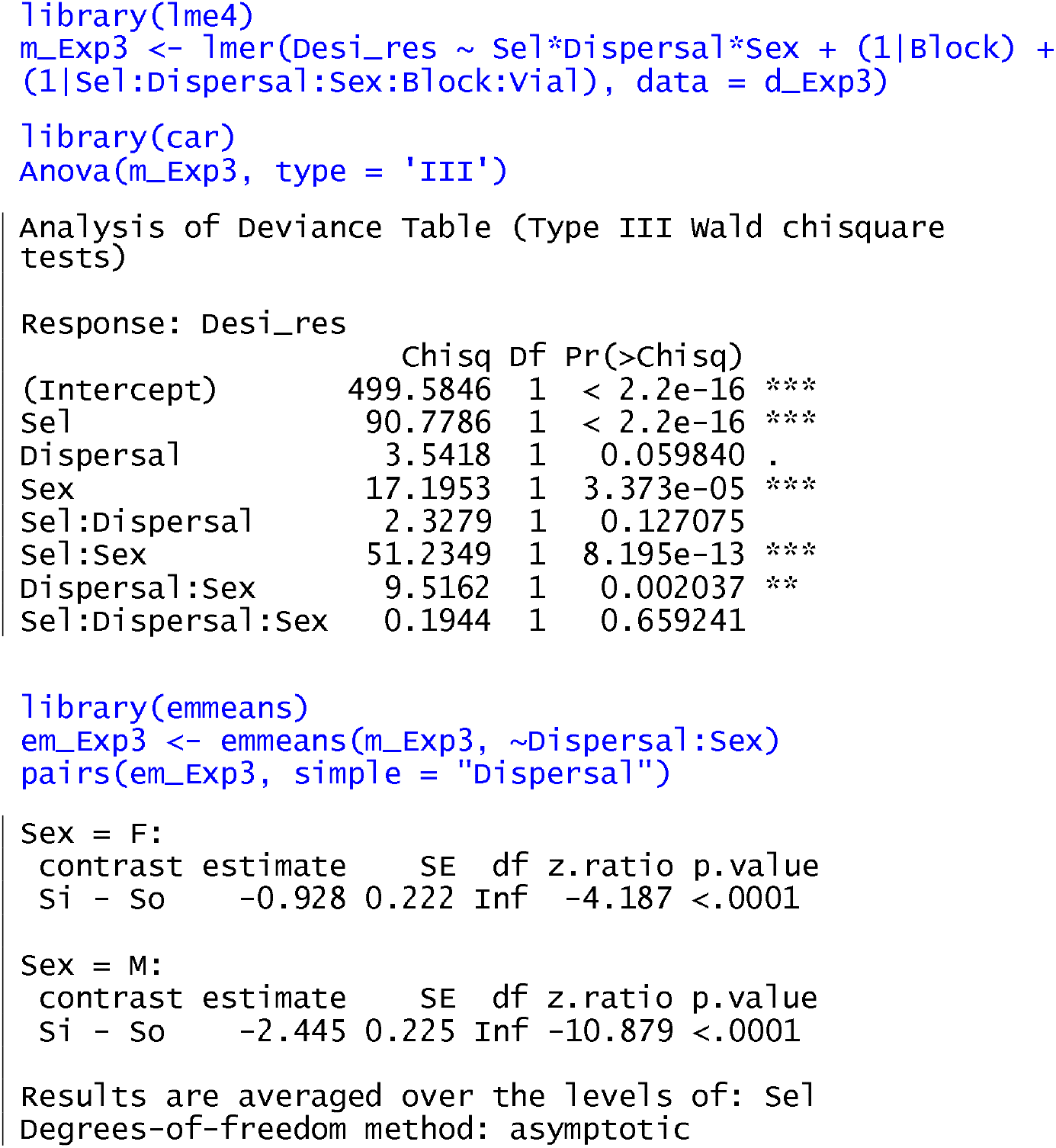

### Text S2.4 Experiment 4

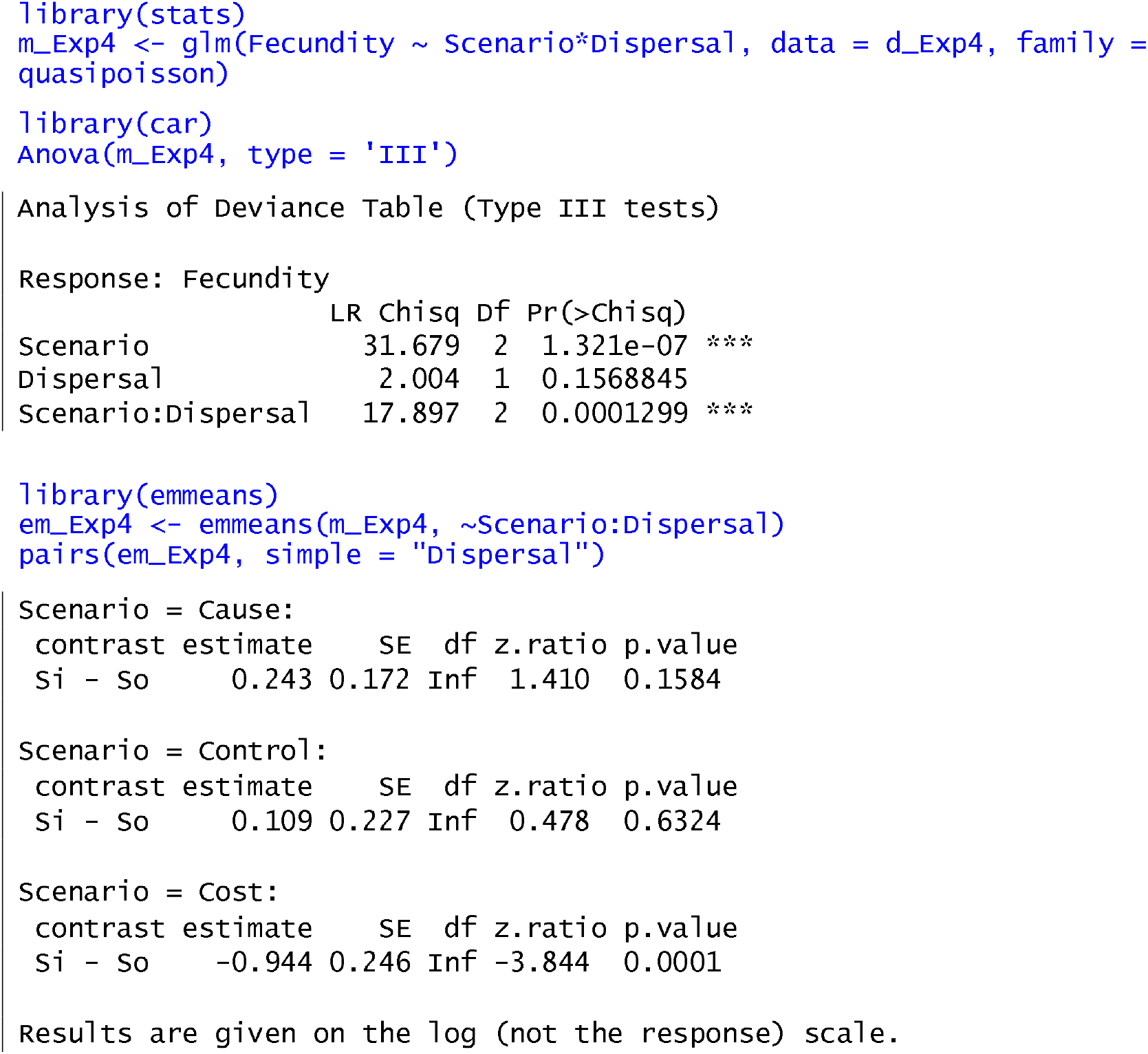

## REFERENCES

Bates, D., Mächler, M., Bolker, B. & Walker, S. (2015) Fitting Linear Mixed-Effects Models Using lme4. Journal of Statistical Software, 67, 1–48.

Bazinet, A.L., Marshall, K.E., MacMillan, H.A., Williams, C.M. & Sinclair, B.J. (2010) Rapid changes in desiccation resistance in Drosophila melanogaster are facilitated by changes in cuticular permeability. Journal of Insect Physiology, 56, 2006–2012.

Black, M. & Pritchard, H.W. (2002) Desiccation and survival in plants: drying without dying. Cabi.

Boeye, J., Travis, J.M.J., Stoks, R. & Bonte, D. (2013) More rapid climate change promotes evolutionary rescue through selection for increased dispersal distance. Evolutionary Applications, 6, 353–364.

Bonte, D. & Dahirel, M. (2017) Dispersal: a central and independent trait in life history. Oikos, 126, 472–479.

Bonte, D., Van Dyck, H., Bullock, J., Coulon, A., Delgado, M., Gibbs, M., Lehouck, V., Matthysen, E., Mustin, K., Saastamoinen, M. & Schtickzelle, N. (2012) Costs of dispersal. Biological Reviews, 87, 290–312.

Bowler, D.E. & Benton, T.G. (2005) Causes and consequences of animal dispersal strategies: relating individual behaviour to spatial dynamics. Biological Reviews, 80, 205–225.

Buckley, L.B., Tewksbury, J.J. & Deutsch, C.A. (2013) Can terrestrial ectotherms escape the heat of climate change by moving? Proceedings of the Royal Society B: Biological Sciences, 280, 20131149.

Cheptou, P.O., Carrue, O., Rouifed, S. & Cantarel, A. (2008) Rapid evolution of seed dispersal in an urban environment in the weed Crepis sancta. Proceedings of the National Academy of Sciences, 105, 3796–3799.

Chippindale, A.K., Ngo, A.L. & Rose, M.R. (2003) The devil in the details of life-history evolution: instability and reversal of genetic correlations during selection onDrosophila development. Journal of genetics, 82, 133–145.

Clark, A. & Doane, W. (1983) Desiccation tolerance of the adipose60 mutant of Drosophila melanogaster. Hereditas, 99, 165–175.

Cohen, J. (1988) Statistical Power Analysis for the Behavioral Sciences. Lawrence Erlbaum Asssociates, USA.

Comte, L. & Olden, J. (2018) Evidence for dispersal syndromes in freshwater fishes. Proceedings of the Royal Society of London, Series B: Biological Sciences, 285, 20172214.

Cremer, S. & Heinze, J. (2003) Stress grows wings: environmental induction of winged dispersal males in Cardiocondyla ants. Current Biology, 13, 219–223.

Djawdan, M., Sugiyama, T.T., Schlaeger, L.K., Bradley, T.J. & Rose, M.R. (2004) Metabolic aspects of the trade-off between fecundity and longevity in Drosophila melanogaster. Methuselah Flies: A Case Study in the Evolution of Aging, pp. 145–164. World Scientific.

Einum, S., SundtCHansen, L. & H. Nislow, K. (2006) The partitioning of densityCdependent dispersal, growth and survival throughout ontogeny in a highly fecund organism. Oikos, 113, 489–496.

Folk, D.G. & Bradley, T.J. (2004) The evolution of recovery from desiccation stress in laboratory-selected populations of Drosophila melanogaster. Journal of Experimental Biology, 207, 2671–2678.

Fox, J. & Weisberg, S. (2019) An R Companion to Applied Regression, Third Edition. Sage, Thousand Oaks CA. https://socialsciences.mcmaster.ca/jfox/Books/Companion/.

Friedenberg, N.A. (2003) Experimental evolution of dispersal in spatiotemporally variable microcosms. Ecology Letters, 6, 953–959.

Gerber, N. & Kokko, H. (2018) Abandoning the ship using sex, dispersal or dormancy: multiple escape routes from challenging conditions. Philosophical Transactions of the Royal Society B: Biological Sciences, 373, 20170424.

Ghalambor, C.K. & Martin, T.E. (2001) Fecundity-survival trade-offs and parental risk-taking in birds. Science, 292, 494–497.

Gibbs, A.G., Chippindale, A.K. & Rose, M.R. (1997) Physiological mechanisms of evolved desiccation resistance in Drosophila melanogaster. Journal of Experimental Biology, 200, 1821–1832.

Graves, J.L., Toolson, E.C., Jeong, C., Vu, L.N. & Rose, M.R. (1992) Desiccation, Flight, Glycogen, and Postponed Senescence in Drosophila metanogaster. Physiological Zoology, 65, 268–286.

Gros, A., Hovestadt, T. & Poethke, H.J. (2008) Evolution of sex-biased dispersal: the role of sex-specific dispersal costs, demographic stochasticity, and inbreeding. Ecological Modelling, 219, 226–233.

Guerra, P. (2011) Evaluating the lifeChistory tradeCoff between dispersal capability and reproduction in wing dimorphic insects: a metaCanalysis. Biological Reviews, 86, 813–835.

Hoffmann, A.A., Hallas, R.J., Dean, J.A. & Schiffer, M. (2003) Low potential for climatic stress adaptation in a rainforest Drosophila species. Science, 301, 100–102.

Hoffmann, A.A. & Harshman, L.G. (1999) Desiccation and starvation resistance in Drosophila: patterns of variation at the species, population and intrapopulation levels. Heredity, 83, 637–643.

Holmstrup, M., Hedlund, K. & Boriss, H. (2002) Drought acclimation and lipid composition in Folsomia candida: implications for cold shock, heat shock and acute desiccation stress. Journal of Insect Physiology, 48, 961–970.

Holzinger, A. & Karsten, U. (2013) Desiccation stress and tolerance in green algae: consequences for ultrastructure, physiological and molecular mechanisms. Frontiers in plant science, 4, 327.

Ives, P.T. (1970) Further genetic studies of the South Amherst population of Drosophila melanogaster. Evolution, 24, 507–518.

Jessup, C. & Bohannan, B. (2008) The shape of an ecological tradeCoff varies with environment. Ecology Letters, 11, 947–959.

Jill, D. & Daniel, R.F. (2003) The effects of size, sex, and reproductive condition on thermal and desiccation stress in a riparian spider (Pirata sedentarius, Araneae, Lycosidae). The Journal of Arachnology, 31, 278–284.

Karan, D. & Parkash, R. (1998) Desiccation tolerance and starvation resistance exhibit opposite latitudinal clines in Indian geographical populations of Drosophila kikkawai. Ecological Entomology, 23, 391–396.

Kellermann, V., Van Heerwaarden, B., Sgrò, C.M. & Hoffmann, A.A. (2009) Fundamental evolutionary limits in ecological traits drive Drosophila species distributions. Science, 325, 1244–1246.

Kranner, I., Beckett, R., Hochman, A. & Nash Iii, T.H. (2008) Desiccation-tolerance in lichens: a review. The Bryologist, 576–593.

Legrand, D., Larranaga, N., Bertrand, R., Ducatez, S., Calvez, O., Stevens, V.M. & Baguette, M. (2016) Evolution of a butterfly dispersal syndrome. Proceedings of the Royal Society of London, Series B: Biological Sciences, 283, 20161533.

Legrand, D., Trochet, A., Moulherat, S., Calvez, O., Stevens, V.M., Ducatez, S., Clobert, J. & Baguette, M. (2015) Ranking the ecological causes of dispersal in a butterfly. Ecography, 38, 822–831.

Lenth, R. (2020) emmeans: Estimated Marginal Means, aka Least-Squares Means. R package version 1.5.2–1. https://CRAN.R-project.org/package=emmeans.

Li, X.Y. & Kokko, H. (2019) Sex-biased dispersal: a review of the theory. Biological Reviews, 94, 721–736.

Lyons, C.L., Coetzee, M., Terblanche, J.S. & Chown, S.L. (2014) Desiccation tolerance as a function of age, sex, humidity and temperature in adults of the African malaria vectors Anopheles arabiensis and Anopheles funestus. Journal of Experimental Biology, 217, 3823–3833.

Maklakov, A.A. & Lummaa, V. (2013) Evolution of sex differences in lifespan and aging: Causes and constraints. Bioessays, 35, 717–724.

Martorell, C. & MartínezCLópez, M. (2014) Informed dispersal in plants: Heterosperma pinnatum (Asteraceae) adjusts its dispersal mode to escape from competition and water stress. Oikos, 123, 225–231.

Matthysen, E. (2012) Multicausality of dispersal: a review. Dispersal ecology and evolution, p. 3–18. Oxford University Press.

Matzkin, L., Watts, T.D. & Markow, T.A. (2007) Desiccation Resistance in Four Drosophila Species: Sex and Population Effects. Fly, 1, 268–273.

Mishra, A., Chakraborty, P.P. & Dey, S. (2020) Dispersal evolution diminishes the negative density dependence in dispersal. Evolution, 74, 2149–2157.

Mishra, A., Tung, S., Shreenidhi, P.M., Sadiq, M.A., Sruti, V.R.S., Chakraborty, P.P. & Dey, S. (2018a) Sex differences in dispersal syndrome are modulated by environment and evolution. Philosophical Transactions of the Royal Society of London, Series B: Biological Sciences, 373, 20170428.

Mishra, A., Tung, S., Sruti, V., Srivathsa, S. & Dey, S. (2020) Mate-finding dispersal reduces local mate limitation and sex bias in dispersal. Journal of Animal Ecology, 89, 2089–2098.

Mishra, A., Tung, S., Sruti, V.R.S., Sadiq, M.A., Srivathsa, S. & Dey, S. (2018b) Pre-dispersal context and presence of opposite sex modulate density dependence and sex bias of dispersal. Oikos, 127, 1596–1604.

Miyatake, T. (1997) Genetic trade-off between early fecundity and longevity in Bactrocera cucurbitae (Diptera: Tephritidae). Heredity, 78, 93–100.

Muller-Landau, H.C. (2010) The tolerance–fecundity trade-off and the maintenance of diversity in seed size. Proceedings of the National Academy of Sciences, 107, 4242–4247.

Parsons, P. (1970) Genetic heterogeneity in natural populations of Drosophila melanogaster for ability to withstand dessication. Theoretical and Applied Genetics, 40, 261–266.

R Core Team (2020) R: A language and environment for statistical comCputing. R Foundation for Statistical Computing, Vienna, Austria. URL https://www.R-project.org.

Rajpurohit, S., Nedved, O. & Gibbs, A.G. (2013) Meta-analysis of geographical clines in desiccation tolerance of Indian drosophilids. Comparative Biochemistry and Physiology Part A: Molecular & Integrative Physiology, 164, 391–398.

Rantala, M.J. & Roff, D.A. (2007) Inbreeding and extreme outbreeding cause sex differences in immune defence and life history traits in Epirrita autumnata. Heredity, 98, 329–336.

Riotte-Lambert, L. & Matthiopoulos, J. (2020) Environmental predictability as a cause and consequence of animal movement. Trends in Ecology & Evolution, 35, 163–174.

Roff, D. (1977) Dispersal in dipterans: its costs and consequences. Journal of Animal Ecology, 46, 443–456.

Roff, D.A. & Fairbairn, D.J. (2007) The Evolution and Genetics of Migration in Insects. Bioscience, 57, 155–164.

Ronce, O. & Clobert, J. (2012) Dispersal syndromes. Dispersal ecology and evolution, pp. 119–138. Oxford University Press.

Sah, P., Salve, J.P. & Dey, S. (2013) Stabilizing biological populations and metapopulations through Adaptive Limiter Control. Journal of Theoretical Biology, 320, 113–123.

Shaw, A.K., Kokko, H. & Neubert, M.G. (2018) Sex difference and Allee effects shape the dynamics of sexCstructured invasions. Journal of Animal Ecology, 87, 36–46.

Stevens, V., Whitmee, S., Galliard, L., Clobert, J., BöhningCGaese, K., Bonte, D., Brändle, M., Matthias Dehling, D., Hof, C., Trochet, A. & Baguette, M. (2014) A comparative analysis of dispersal syndromes in terrestrial and semiCterrestrial animals. Ecology Letters, 17, 1039–1052.

Tigreros, N. & Davidowitz, G. (2019) Flight-fecundity tradeoffs in wing-monomorphic insects.

Travis, J., Delgado, M., Bocedi, G., Baguette, M., Bartoń, K., Bonte, D., Boulangeat, I., Hodgson, J., Kubisch, A., Penteriani, V. & Saastamoinen, M. (2013) Dispersal and species’ responses to climate change. Oikos, 122, 1532–1540.

Trochet, A., Courtois, E.A., Stevens, V.M., Baguette, M., Chaine, A., Schmeller, D.S., Clobert, J. & Wiens, J.J. (2016) Evolution of sex-biased dispersal. The Quarterly Review of Biology, 91, 297–320.

Tuba, Z., Slack, N.G. & Stark, L.R. (2011) Bryophyte ecology and climate change. Cambridge University Press.

Tung, S., Mishra, A., Gogna, N., Sadiq, M.A., Shreenidhi, P.M., Sruti, V.R.S., Dorai, K. & Dey, S. (2018a) Evolution of dispersal syndrome and its corresponding metabolomic changes. Evolution, 72, 1890–1903.

Tung, S., Mishra, A., Shreenidhi, P.M., Sadiq, M.A., Joshi, S., Sruti, V.R.S. & Dey, S. (2018b) Simultaneous evolution of multiple dispersal components and kernel. Oikos, 127, 34–44.

Van Heerwaarden, B. & Sgrò, C.M. (2014) Is adaptation to climate change really constrained in niche specialists? Proceedings of the Royal Society B: Biological Sciences, 281, 20140396.

Wilkin, T.A. & Sheldon, B.C. (2009) Sex Differences in the Persistence of Natal Environmental Effects on Life Histories. Current Biology, 19, 1998–2002.

